# A machine learning approach to predicting autism risk genes: Validation of known genes and discovery of new candidates

**DOI:** 10.1101/463547

**Authors:** Ying Lin, Anjali M. Rajadhyaksha, James B. Potash, Shizhong Han

## Abstract

Autism spectrum disorder (ASD) is a complex neurodevelopmental condition with a strong genetic basis. The role of *de novo* mutations in ASD has been well established, but the set of genes implicated to date is still far from complete. The current study employs a machine learning-based approach to predict ASD risk genes using features from spatiotemporal gene expression patterns in human brain, gene-level constraint metrics, and other gene variation features. The genes identified through our prediction model were enriched for independent sets of ASD risk genes, and tended to be differentially expressed in ASD brains, especially in the frontal and parietal cortex. The highest-ranked genes not only included those with strong prior evidence for involvement in ASD (for example, *TCF20* and *FBOX11*), but also indicated potentially novel candidates, such as *DOCK3*, *MYCBP2* and *CAND1*, which are all involved in neuronal development. Through extensive validations, we also showed that our method outperformed state-of-the-art scoring systems for ranking ASD candidate genes. Gene ontology enrichment analysis of our predicted risk genes revealed biological processes clearly relevant to ASD, including neuronal signaling, neurogenesis, and chromatin remodeling, but also highlighted other potential mechanisms that might underlie ASD, such as regulation of RNA alternative splicing and ubiquitination pathway related to protein degradation. Our study demonstrates that human brain spatiotemporal gene expression patterns and gene-level constraint metrics can help predict ASD risk genes. Our gene ranking system provides a useful resource for prioritizing ASD candidate genes.

## Introduction

Autism spectrum disorder (ASD) is a neurodevelopmental condition characterized by impaired social interaction and communication, as well as repetitive behavior. While its etiology is complex, ASD has a strong genetic basis (1-3). The role of *de novo* mutations in ASD has been firmly established through candidate gene (4, 5), whole exome (6-9), and whole genome sequencing studies (10, 11). Although the list of risk genes implicated by *de novo* mutations is growing, it is still very likely far from complete, with an estimated full set of ASD genes ranging from several hundred to more than 1,000 (9). In the search for additional *de novo* mutations, sequencing studies continue to be an important approach, but the current sequencing cost is still very high, especially for large samples. As an alternative strategy, advanced analytical approaches, which leverage previously implicated genes and prior knowledge, have the potential to enhance risk gene discovery in an efficient and cost-effective manner.

One approach is based on the concept of guilt-by-association, i.e., assuming that genes that confer risk for ASD are likely to be functionally related, and that they thus converge on molecular networks and biological pathways implicated in disease (12, 13). For example, one study showed that ASD genes with *de novo* mutations converged on pathways related to chromatin remodeling and synaptic function (14). To leverage these functional relationships, several studies have explored integrating known risk genes using a protein-protein interaction (PPI) network to identify novel genes involved in ASD (15-18). However, a PPI network is built upon general protein-protein interactions without reference to tissue or cell type specificity, and this approach may not fully capture the brain-centric functional relationships among ASD genes. Accordingly, a brain-specific network-based approach, which considered relationships within the context of the brain, was proposed to predict ASD genes, (19, 20). Studies employing this paradigm, however, did not consider the dynamic patterns of gene relationships during brain development, thereby limiting their potential for discovery, given the possibility that genes might only be functionally related within a specific developmental stage. Evidence for this comes from Willsey et al., who showed, using spatiotemporal gene expression data from human brain, that co-expression patterns of ASD risk genes varied by spatiotemporal windows, with the strongest co-expression patterns observed in the prefrontal and primary motor–somatosensory cortical regions during midfetal development, suggesting an important convergence of risk gene activity in particular places at a particular time (21).

In addition to having functional relationships, ASD genes affected by *de novo* mutations tend to be intolerant of variations (22, 23). With the availability of sequencing data from large samples, recent work has developed measures to quantify the sensitivity of genes to disruptive functional variations (24, 25). Utilizing exome data on more than 60,000 individuals from the Exome Aggregation Consortium (ExAC), a gene-level constraint metric---the probability of being loss-of-function (LoF) intolerant (pLI)---was created, which separates genes into LoF intolerant or LoF tolerant (25). Kosmicki et al. further demonstrated that the excess of *de novo* mutations in ASD individuals was primarily driven by LoF-intolerant genes, but not LoF-tolerant genes (26). To our knowledge, there have been no studies that have quantitatively examined the potential of using gene-level constraint metrics to predict ASD genes.

We reasoned that ASD risk genes show expression patterns that are clustered in specific brain regions and developmental stages critical to disease development, and that high resolution spatiotemporal gene expression patterns in human brain can help distinguish genes that cause disease from those that do not. In addition, because ASD genes affected by *de novo* mutations are sensitive to mutational changes, we reasoned that gene-level constraint metrics can further differentiate ASD genes from normal ones. The objective of this study was to employ a machine learning-based approach to predict ASD risk genes using human brain spatiotemporal gene expression signatures, gene-level constraint metrics, and other gene variation features. We compared the performance of our method with five other state-of-the-art scoring systems for ranking ASD candidate genes, and evaluated the risk genes from our prediction model using independent sets of risk genes and differential gene expression evidence. Gene Ontology (GO) enrichment analysis was also performed to understand the biology underlying ASD risk genes.

## Methods

### Gene set

To train the gene prediction model, we used labeled genes curated by Duda et al, as described in detail elsewhere (20). Briefly, the labeled genes contained 143 true positive genes and 1,145 true negative ones. The true positives came from the high confidence genes in the Simons Foundation Autism Research Initiative (SFARI) resource (https://www.sfari.org/resource/sfari-gene/, Category 1, Category 2, and syndromic genes) and the 65 reported genes in Sanders et al. (27). The true negative genes were selected from the non-ASD gene list created by Krishnan et al. (19), which were genes associated with non-mental health diseases, as annotated in OMIM. Among these genes we focused on those that had both gene expression data from the BrainSpan atlas and gene-level constraint metrics available, so that our final training gene set included 121 true positive genes and 963 true negatives.

### Spatiotemporal gene expression

We downloaded RNA-Seq data (version 10), summarized to Gencode v20 gene-level reads per kilobase million mapped reads (RPKM) values, from the BrainSpan website (http://www.brainspan.org/). Detailed information on tissue processing, experimental and bioinformatics procedures related to the RNA-Seq data is available at the BrainSpan website. The BrainSpan dataset includes 524 gene-level expression features across 13 developmental stages in 31 brain regions. Gene expression values were log-transformed (log_2_ [RPKM□+□1]) and were used to predict autism genes.

To capture the functional relationships among genes, we built a weighted network for genes with both gene co-expression and PPI evidence from InWeb (28). Specifically, the co-expression level between a gene pair was assessed by the Fischer z-transformed Pearson correlation between their spatiotemporal gene expression values. The genes with protein-protein interactions were connected and their edges were weighted by their co-expression levels. We extracted a set of network features that characterized the network topologies using *igraph* package in R. Specifically, we measured the node centralities using node degrees, clones centralities, betweenness centralities, Boncich power centralities, eigenvector centralities, and alpha centralities (29). We captured the modules in functional relationship networks using the principle component decomposition and K-core decomposition (30). The loading of the 1^st^ principle component, hub score and coreness were obtained for each node. The importance of each node was further measured using the PageRank algorithm (31), which counts the number and weight of links to each node. In total, 13 network features were extracted from the weighted gene network and were used for autism risk gene prediction.

### Gene-level constraint metrics and other gene variation features

We used gene-level constraint metrics developed from the exome data of more than 60,000 individuals from the Exome Aggregation Consortium to quantify the sensitivity of genes to variations (25). We considered six gene-level constraint metrics, including Z scores for synonymous (syn_z), missense (mis_z), and LoF variants (lof_z), the probability of being LoF intolerant (pLI), the probability of being intolerant of homozygous but not heterozygous LoF variants (pRec), and the probability of being tolerant of both heterozygous and homozygous LoF variants (pNull). A higher Z score or pLI indicates that the gene is more intolerant of variation (more constrained). We also included 10 general gene features, including the number of coding base pairs (bp), probabilities of mutations across the transcript for synonymous (mu_syn), missense (mu_mis), and LoF variants (mu_lof), number of rare variants (n_syn, n_mis, n_lof), and depth adjusted number of expected rare variants (exp_syn, exp_mis, exp_lof). Wilcoxon rank sum test was used to compare the group differences in above features between known ASD risk and non-risk genes.

### Autism risk gene prediction

We used machine learning methods to predict autism risk genes from their spatiotemporal expression signatures, network topology features, gene-level constraint metrics, and other general gene features. We applied four machine learning methods ranging from ones that are regression based [logistic regression and support vector machines (SVM) with Gaussian kernel] to others that are tree based (random forest and boosted trees). The optimal tuning parameters in each model were selected by a nested grid-search, and model performances were evaluated by five-fold cross validation (CV) on training data. The prediction accuracy was measured by the receiver-operator (ROC) curve and the area under the receiver-operator curve (AUC) on the hold-out set for each fold of the CV.

Based on the average prediction accuracy over five folds, the random forest model was selected as the optimal algorithm, and the corresponding prediction models were stored to predict over 17,000 unlabeled genes, which had both spatial-temporal expression signatures and constraint metrics available. Each gene was assigned a risk score indicating its likelihood of being involved in autism. The risk score was predicted as the average of scores from the five prediction models that emerged from each CV. For each labeled gene, the risk score was obtained by the prediction model that left the gene held out of the set in the CV.

### Autism risk gene validation using differential gene expression evidence

Based on our gene ranking system, we classified genes into risk and non-risk genes using a threshold of risk score of 0.2, which gave the highest prediction accuracy on training data. Genes with a risk score higher than the threshold were predicted as risk genes and the remaining genes were predicted as non-risk genes. We validated the classification performance by examining whether our predicted risk genes show differential gene expression evidence for ASD. Specifically, we obtained differential gene expression summary statistics (beta and p-values) for ASD from RNA-Seq datasets for four major cortical lobes (frontal, temporal, parietal, and occipital) and their average, as well as the summary statistics for inflammatory bowel disease (IBD), a non-psychiatric disorder that we employed as a negative control (32). We used simulation-based approach to estimate the enrichment statistics of predicted risk genes in differential gene expression evidence. We first generated a background distribution from random gene sets, while matching for gene size found in predicted risk genes. The enrichment fold was estimated by the ratio of the observed number of risk genes with differential gene expression evidence (p < 0.05) to the average number of that from random gene sets. The p-value for enrichment was then the proportion of random gene sets with the same or a greater number of genes with differential gene expression evidence, as compared to the number found for predicted risk genes. To investigate whether the enrichment of differential gene expression evidence was specific to ASD, we also performed the same enrichment analysis for IBD.

### Autism risk gene validation in independent sequencing studies

We further evaluated our gene ranking system utilizing genes targeted by *de novo* LoF mutations from two independent studies, including one that performed whole exome sequencing of 2,517 families in the Simons Simplex Collection (SSC) cohort (9) and another that performed whole genome sequencing of the MSSNG cohort (10). For the SSC cohort, after excluding genes not included in BrainSpan, we compiled a list of 27 recurrent LoF *de novo* mutations in probands, 346 singleton LoF *de novo* mutations in probands, and 170 LoF *de novo* mutations in the unaffected siblings as negative controls. From the study of the MSSNG cohort, we created a list of 212 *de novo* LoF mutations in probands, 58 statistically significant *de novo* LoF or missense mutations, and 18 statistically significant *de novo* LoF or missense mutations that were not previously reported. For each of the six gene lists, we tested whether a larger proportion of genes were observed in the first decile of our gene ranking system than expected using a binomial test. The expected proportion (0.166) was determined using the percentage of genes with synonymous *de novo* mutations in the unaffected siblings of the SSC cohort.

### Gene Ontology enrichment analysis

We performed Gene Ontology (GO) enrichment analysis to examine whether predicted risk genes were clustered into specific biological processes. Fisher’s exact test was used to test the enrichment of risk genes in GO terms compared to non-risk genes. GO terms were chosen from the GO ontology of biological processes in MSigDB (v5.2) (33). To facilitate interpretation of the results, we included 2,758 GO terms that overlapped at least 20, but not more than 2,000 genes with our tested genes. Bonferroni correction was applied for multiple testing correction. Because GO terms were often highly overlapping in genes, we used hierarchical clustering to group significant gene sets into clusters based on similarity of their gene profiles (34). We first defined a gene overlapping matrix by counting the number of overlapping genes for each pair of gene sets. The Pearson correlation coefficient *R* was then calculated for each pair of gene sets based on their overlap profiles. The distance matrix for hierarchical clustering was then 1 − *R*. Hierarchical clustering was performed using the “ward” method implemented in the R function “hclust”. The dendrogram and heatmap were plotted using the R function “heatmap.2”.

### Comparison with other ranking systems

We compared our predictions with five third-party autism gene prediction scores recently added to the SFARI Gene database, including the ExAC score (pLI) (25), Krishman probability score (19), Iossifov probability score (23), Sanders TADA score (27), and Zhang D score (35). Different gene scoring systems were compared in terms of labeled genes (true positive genes and true negative genes) and unlabeled genes (curated candidate genes) separately. For the labeled genes, the prediction accuracy of gene scores was assessed using the ROC curves and the area under ROC curves. To compare the scores on unlabeled genes, we curated an independent set of 173 autism candidate risk genes, including 130 genes with suggestive evidence from the SFARI Gene database (Category 3) and 43 recurrent *de novo* LoF genes discovered in recent studies (4, 5, 10, 36). We compared the overall ranking of candidate risk genes for different gene scoring systems, with a higher ranking (smaller number) indicating a greater likelihood of being ASD risk genes. We also compared the enrichment of candidate genes in the first decile of different gene scoring systems.

## Results

### Genome-wide prediction of autism risk genes

We visualized gene expression patterns for 1,084 training genes across various regions and developmental stages of human brain (Supplementary Figure 1). There was a trend for known autism risk genes (left gene panel, red rows) to have higher expression levels than non-risk genes (left gene panel, blue rows). We further tested expression level differences between known risk and non-risk genes for each specific brain region and developmental stage (Figure 1). The known autism risk genes showed significantly higher expression levels on average than non-risk genes for all tested brain regions and developmental stages (p < 0.05). Of note, the difference was stronger for early-to-middle prenatal stages, ranging from 12 to 21 post-conceptional weeks (pcw).

**Figure 1.**
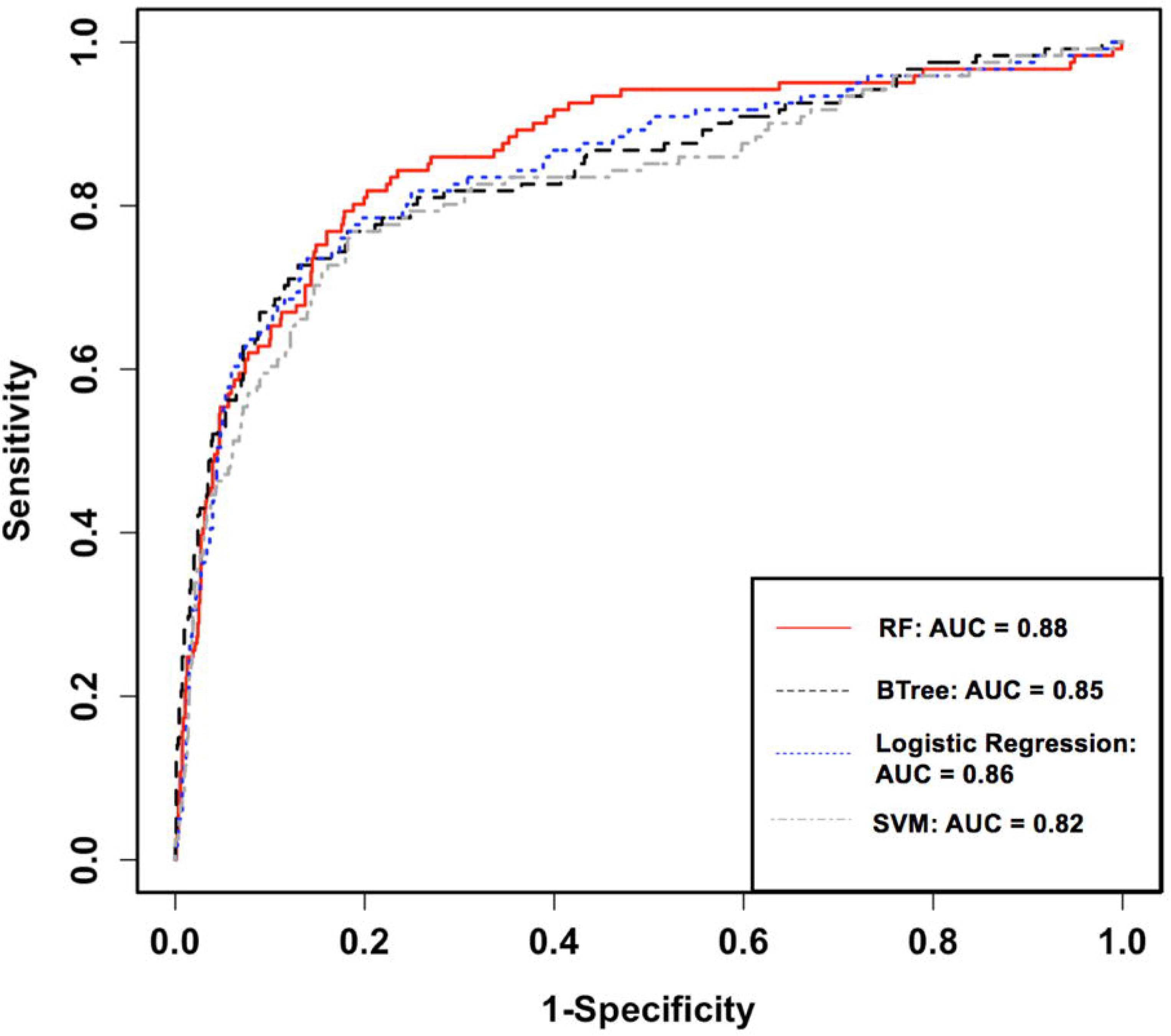
Gene expression difference between ASD risk and non-risk genes in the spatiotemporal development of human brain. Each cell in the heat map represents the expression level difference (t-test) in a specific brain region (column) and development stage (row). The intensity of color represents the log-transformed p-value from a t-test. The brain regions and stages without gene expression data are marked as black.

We compared known autism risk and non-risk genes in their sensitivity to mutational changes and other gene features. As shown in Supplementary Figure 2, compared to non-risk genes, autism risk genes were more intolerant of missense (mis_z, p = 7×10^−16^) and LoF mutations (lof_z, p = 2×10^−23^; pLI, p = 2×10^−22^), were less likely intolerant of homozygous, but not heterozygous LoF variants (pRec, p = 5×10^−21^), and had a lower probability of being tolerant of both heterozygous and homozygous LoF variants (pNull, p = 3×10^−24^). Autism risk genes had longer coding base pairs (p = 4×10^−29^), a higher probability of mutation across the transcript (mu_syn, p = 1×10^−16^; mu_mis, p = 2×10^−18^; mu_lof, p = 4×10^−19^), and a larger number of rare synonymous or missense variants (n_syn, p = 4×10^−16^; n_mis, p = 1×10^−6^).

We compared the prediction accuracy of four machine learning algorithms using five-fold CV. The random forest model achieved the best prediction accuracy for autism risk genes with an AUC of 0.88 (Figure 2). The effects of different features on the random forest model were further explored by comparing the prediction accuracy under different feature sets (Supplementary Figure 3). We found that using the spatiotemporal gene expression features alone achieved an AUC greater than 0.8, and that the prediction accuracy was further improved by including gene network features, gene-level constraint metrics, and other gene features. We further evaluated the importance of individual features in the optimal random forest model. The feature importance was quantified as the average gain, i.e., improvement in node purity, of the feature when it was used in trees. Figure 3 illustrates the top 30 important features, including 14 spatiotemporal expression features, two gene network features, six gene-level constraint metrics, and eight other gene features.

**Figure 2.**
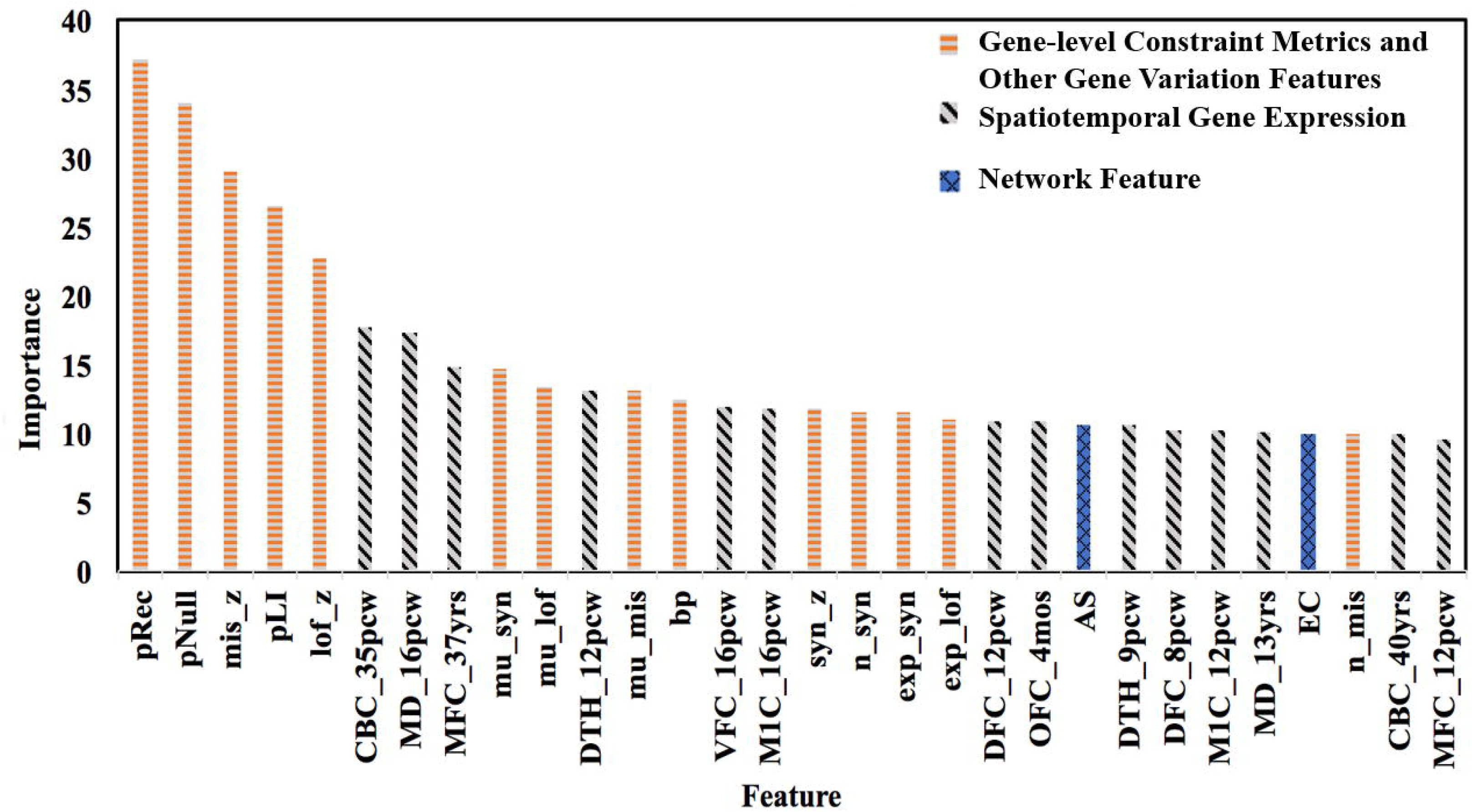
Performance of four machine learning algorithms in five-fold Cross Validation. The performance was measured by ROC curve and the area under curve (AUC).

**Figure 3.**
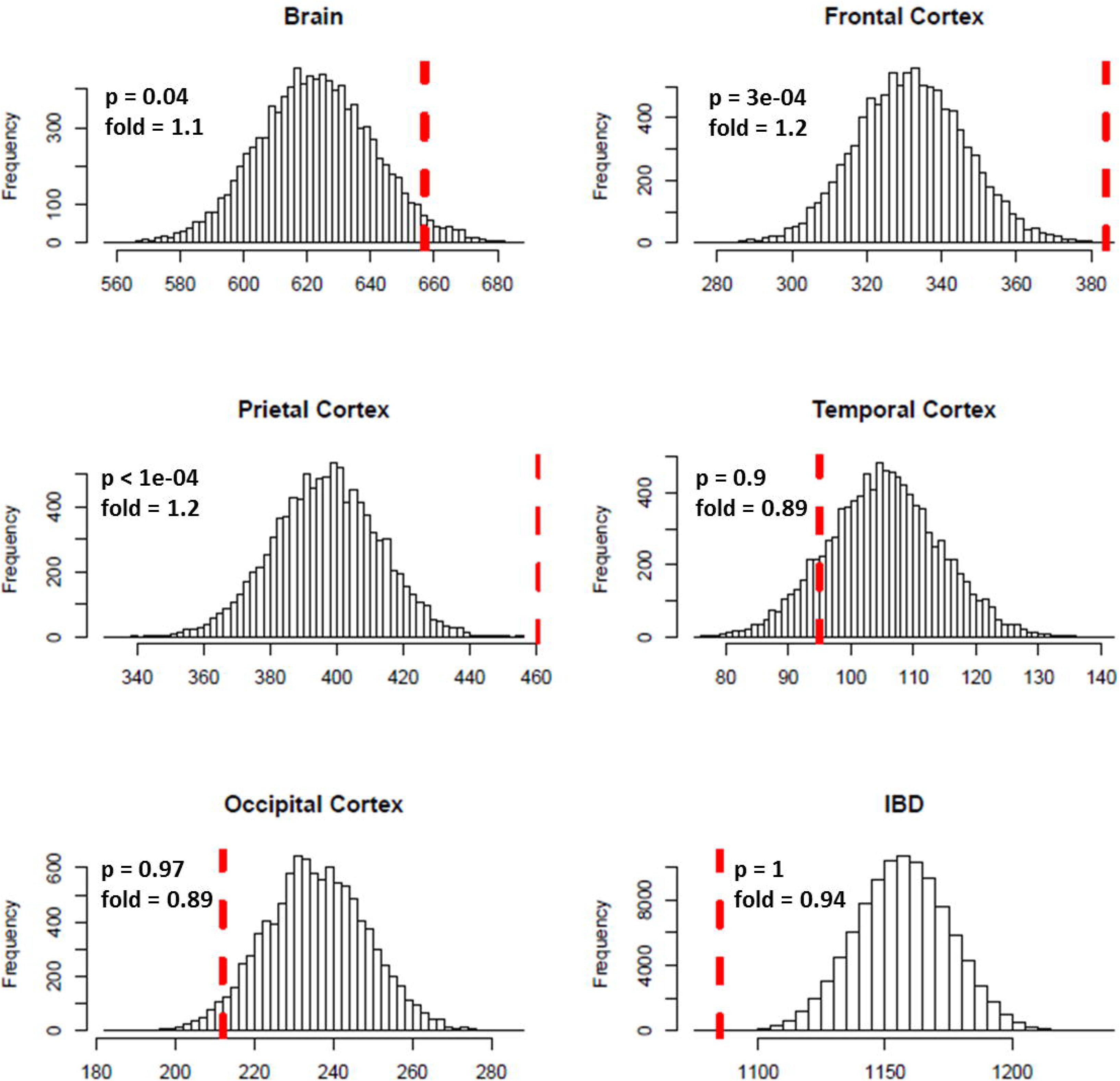
Top 30 important features in the random forest model.

### Autism risk gene validation using differential gene expression evidence

We predicted 2,098 risk genes using our gene ranking system. We then examined whether those predicted risk genes were enriched for differential gene expression evidence for ASD. We found that the predicted risk genes tended to be more differentially expressed in ASD brains, especially in frontal (fold = 1.2, p = 3.0 × 10^−4^) and parietal cortex (fold = 1.2, p < 1.0 × 10^−4^) (Figure 4). Interestingly, we found that the enrichment was primarily driven by down regulated genes in autism brain (frontal cortex, fold = 1.6, p < 1.0 × 10^−4^; parietal cortex, fold = 1.7, p < 1.0 × 10^−4^) (Supplementary Figure 4). We did not see any significant enrichment of differential gene expression evidence for IBD, suggesting that the enriched DGE in our predicted genes was specific to ASD.

**Figure 4.**
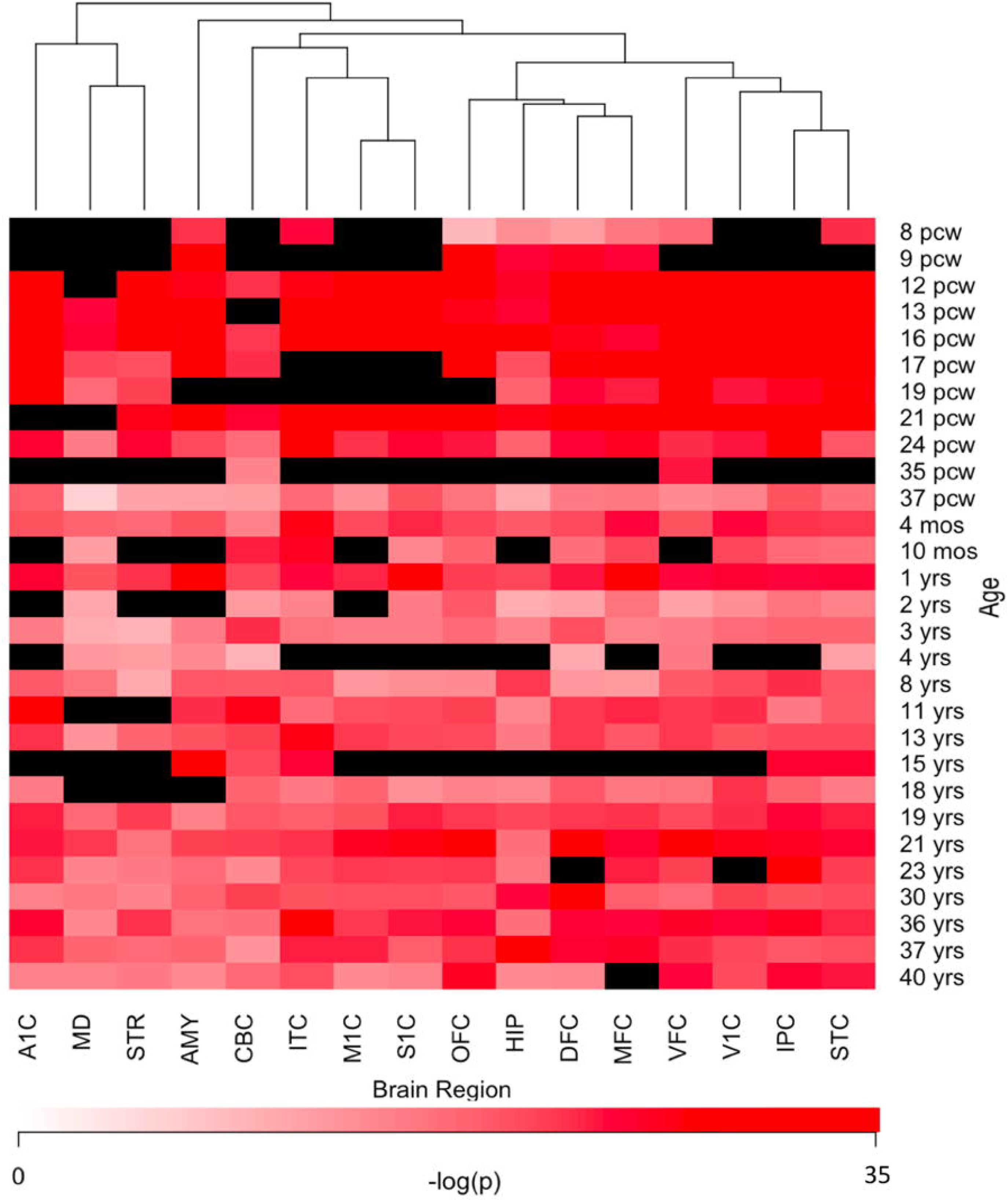
Enrichment analysis of differential expression evidence for predicted ASD risk genes.

### Autism risk gene validation in independent sequencing studies

We further evaluated our gene ranking system using two independent sequencing studies (Figure 5). For the risk genes identified from the SSC cohort, our top decile genes were significantly enriched with *de novo* LoF mutations in probands. Specifically, genes in the first decile of our ranking system included 89% (25 of 27, p = 7.8 × 10^−18^) of the recurrent *de novo* LOF mutations and 35% (121 of 346, p = 9.9 × 10^−17^) of *de novo* LOF mutations in probands. In contrast, we did not observe significant enrichment of genes with *de novo* LOF mutations in the unaffected siblings (p = 0.55). Similarly, for risk genes identified from the MSSNG cohort, we found significant enrichment for all three gene lists, including the *de novo* LOF mutations in probands (28%, p = 2.8 × 10^−5^), the 58 genes that reached genome-wide significance (74%, p = 6.0 × 10^−22^), and the 18 novel genes (56%, p = 0.0002).

**Figure 5.**
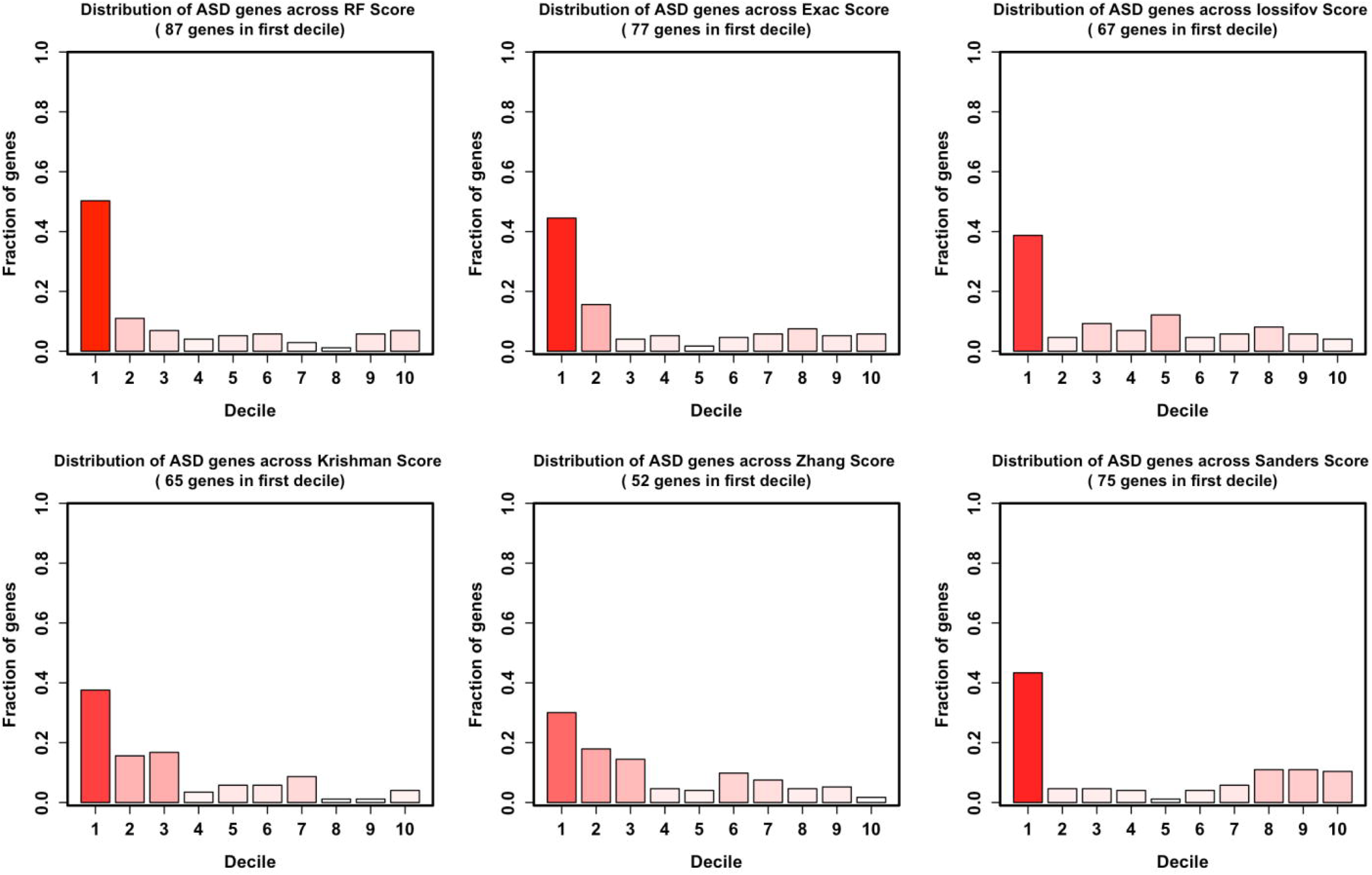
Decile enrichment of ASD-related genes from two independent cohorts in our gene ranking system.

### Gene Ontology enrichment analysis

We conducted GO enrichment analysis to examine whether predicted risk genes were clustered into specific biological processes. The full results of this analysis are shown in Supplementary Table 1. There were 276 GO terms that remained significant after Bonferroni correction (p_corrected_ < 0.05). Significant GO terms were grouped into five major clusters using hierarchical clustering (Supplementary Figure 5). These clusters included GO terms related to neuronal signaling (black), neurogenesis (red), chromatin remodeling (blue), transcriptional regulation (yellow), and protein processing (orange). Table 2 shows details for the top 10 enriched GO terms that were particularly interesting, as they included GO terms involved in ionotropic glutamate receptor signaling, motor neuron axon guidance, regulation of RNA alternative splicing, and protein dealkylation.

**Table 2.** Top ten enriched GO terms in predicted ASD risk genes.

| **GO terms** | **OR** | **95%_CI_L** | **95%_CI_U** | **p<sub>adj</sub>** | **cluster** |
| --- | --- | --- | --- | --- | --- |
| GO_IONOTROPIC_Glutamate_Receptor_Signaling_Pathway | 24.5 | 8.7 | 85.0 | 1.21E-08 | black |
| GO_Central_Nervous_System_Projection_Neuron_Axonogenesis | 21.6 | 7.5 | 76.1 | 4.80E-07 | red |
| GO_Central_Nervous_System_Neuron_Axonogenesis | 14.4 | 5.8 | 38.9 | 1.92E-06 | red |
| GO_Positive_Regulation_of_RNA_Splicing | 13.5 | 5.4 | 36.8 | 1.06E-05 | yellow |
| GO_Dendrite_Morphogenesis | 13.2 | 6.3 | 29.4 | 2.81E-09 | red |
| GO_Glutamate_Receptor_Signaling_Pathway | 11.9 | 5.8 | 25.0 | 2.96E-09 | black |
| GO_Motor_Neuron_Axon_Guidance | 11.2 | 4.5 | 29.4 | 0.000128 | red |
| GO_Dendrite_Development | 9.7 | 5.7 | 16.6 | 5.59E-14 | red |
| GO_Modulation_of_Excitatory_Postsynaptic_Potential | 9.4 | 3.8 | 23.8 | 0.001314 | black |
| GO_Protein_Dealkylation | 9.2 | 3.9 | 22.3 | 0.000551 | blue |

OR: odds ratio; 95%\_CI\_L: OR 95% confidence interval lower bound; 95%\_CI\_U: OR 95% confidence interval upper bound

### Comparison with other ranking systems

We compared the performance of our ranking system (RF, random forest) with five other gene scoring systems in their ability to rank labeled or unlabeled genes. Our ranking system outperformed the other scores on classifying labeled genes, as it displayed the highest AUC value 0.88 (Supplementary Figure 6). When we examined the rank of an independent set of 173 autism candidate genes, our method again outperformed other methods, because our method had the smallest median ranking (indicating the greatest likelihood of the set containing autism risk genes) (Supplementary Figure 7). We further compared the enrichment of 173 candidate genes in the first decile of each gene ranking system (Figure 6). We observed the highest proportion of candidate genes in the first decile of our ranking system (50%), which was higher than the ExAC score (45%), Sanders TADA score (43%), Iossifov probability score (39%), Krishman probability score (38%), and Zhang D score (30%). In summary, using both labeled and unlabeled genes, we showed that our method improved the performance of ranking ASD candidate genes.

**Figure 6.**
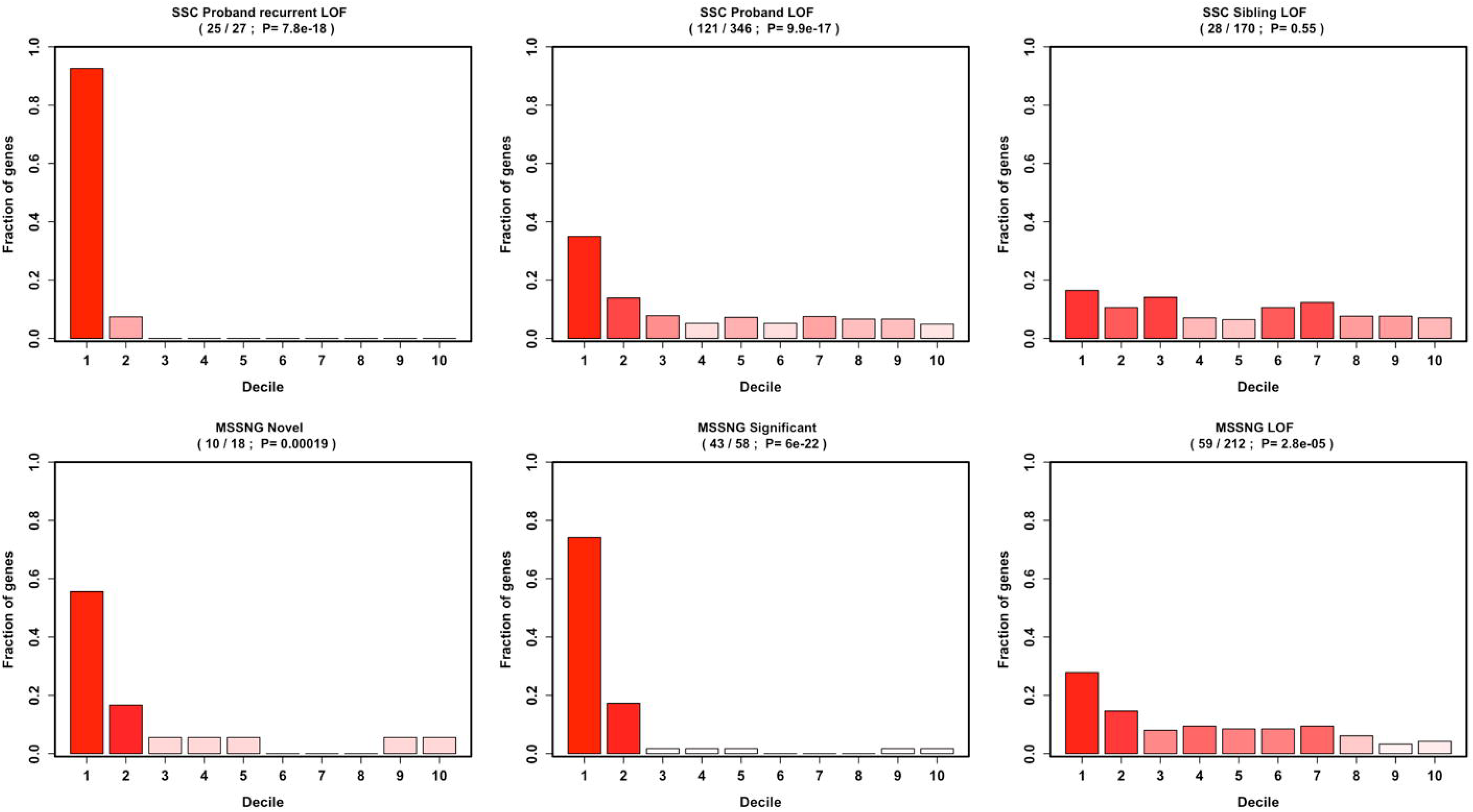
Comparison of the proportion of 173 independent candidate genes in the first decile of each gene ranking system.

### The role of highest-ranked genes in ASD

We examined the function and the role of our highest-ranked genes (top ten) in ASD (table 1). Among the top ten ranked genes, five have been reported to be involved in ASD (*HERC1*, *NBEA*, *ATRX*, *FBXO11*, and *TCF20*). We did not find previous direct association evidence with ASD for the other five genes (*CAND1*, *ZYG11B*, *DOCK3*, *XPO7*, and *MYCBP2*), suggesting these are potentially novel candidates. Three of these genes in particular stood out (*DOCK3*, *MYCBP2,* and *CAND1*) for their known involvement in neuronal development. In addition, among the top ten ranked genes, five were related to the ubiquitination pathway (*HERC1, CAND1, ZYG11B, FBXO11,* and *MYCBP2*), indicating the role of protein ubiquitination pathway in ASD. Interestingly, all, but three of top ten ranked genes show reduced gene expression in ASD across four cortical lobes (p < 0.05) (Supplementary Table 3). The strongest differential gene expression evidence was observed for *DOCK3* (p=0.00025). The lack of difference in gene expression of the three genes may be due to their lower expression postnatally, especially for *TCF20* (Supplementary Figure 8).

**Table 1.**
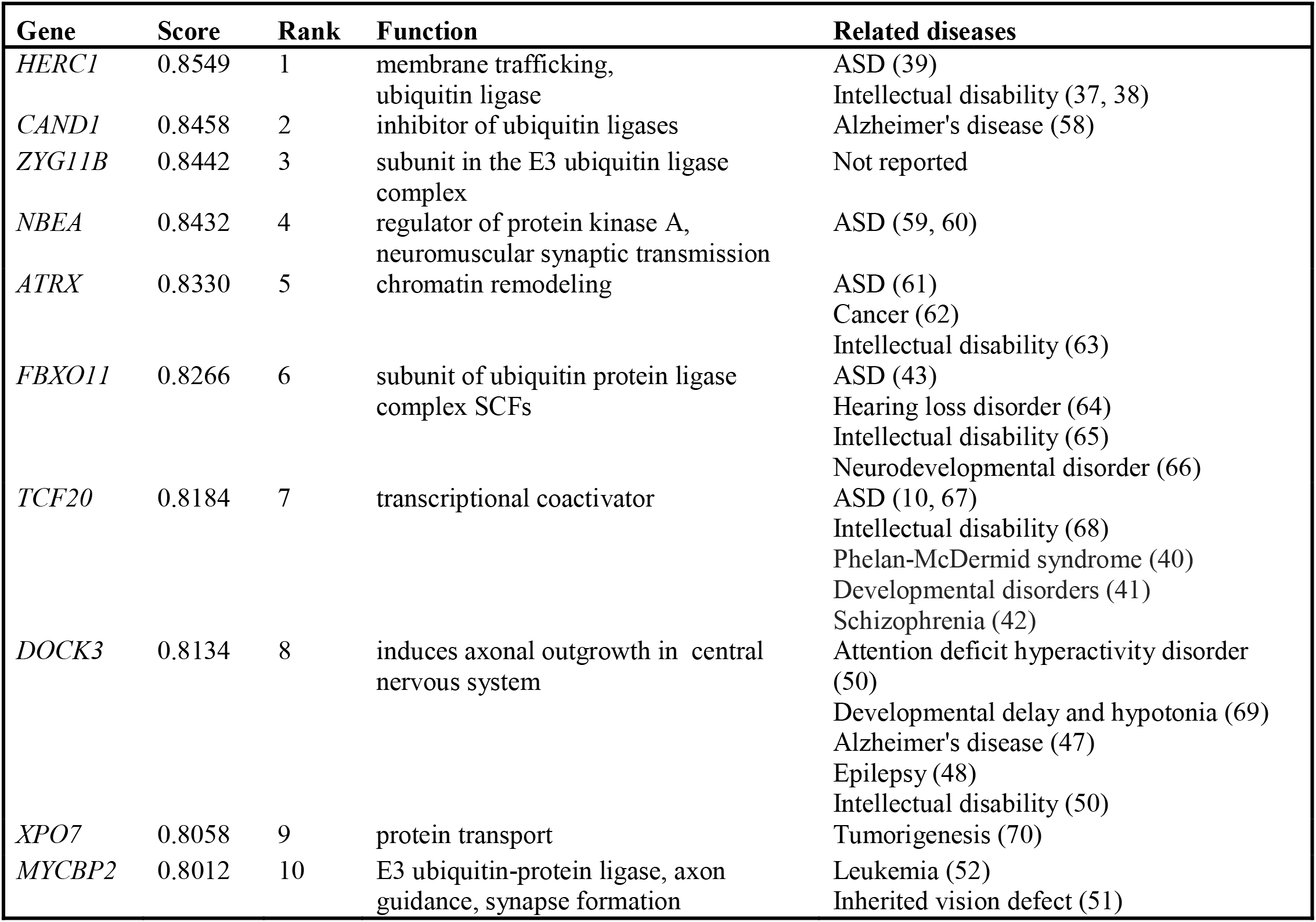
Function and roles of our highest-ranked genes in ASD and other diseases.

| **Gene** | **Score** | **Rank** | **Function** | **Related diseases** |
| --- | --- | --- | --- | --- |
| *HERC1* | 0.8549 | 1 | membrane trafficking, ubiquitin ligase | ASD (39)<br>Intellectual disability (37, 38) |
| *CAND1* | 0.8458 | 2 | inhibitor of ubiquitin ligases | Alzheimer's disease (58) |
| *ZYG11B* | 0.8442 | 3 | subunit in the E3 ubiquitin ligase complex | Not reported |
| *NBEA* | 0.8432 | 4 | regulator of protein kinase A, neuromuscular synaptic transmission | ASD (59, 60) |
| *ATRX* | 0.8330 | 5 | chromatin remodeling | ASD (61)<br>Cancer (62)<br>Intellectual disability (63) |
| *FBXO11* | 0.8266 | 6 | subunit of ubiquitin protein ligase complex SCFs | ASD (43)<br>Hearing loss disorder (64)<br>Intellectual disability (65)<br>Neurodevelopmental disorder (66) |
| *TCF20* | 0.8184 | 7 | transcriptional coactivator | ASD (10, 67)<br>Intellectual disability (68)<br>Phelan-McDermid syndrome (40)<br>Developmental disorders (41)<br>Schizophrenia (42) |
| *DOCK3* | 0.8134 | 8 | induces axonal outgrowth in central nervous system | Attention deficit hyperactivity disorder (50)<br>Developmental delay and hypotonia (69)<br>Alzheimer's disease (47)<br>Epilepsy (48)<br>Intellectual disability (50) |
| *XPO7* | 0.8058 | 9 | protein transport | Tumorigenesis (70) |
| *MYCBP2* | 0.8012 | 10 | E3 ubiquitin-protein ligase, axon guidance, synapse formation | Leukemia (52)<br>Inherited vision defect (51) |

## Discussion

A number of methods have been developed for inferring ASD risk genes. Although they employ differing computational methodologies, conceptually, most methods were built upon the concept of guilt-by-association, using the assumption that risk genes are functionally related. Theoretically, ASD risk genes should exert their effects at specific developmental stages in specific brain tissues or cell types that are critical to disease development. However, most existing methods have not considered the spatial and temporal patterns of gene relationships during brain development. In addition, gene-level constraint metrics, such as loss of function intolerance, have been used to prioritize ASD candidate genes, but no studies have quantitatively examined their potential for predicting ASD genes. To our knowledge, the current study represents the first method that explicitly uses human brain spatiotemporal gene expression features and gene-level constraint metrics for predicting ASD genes. Using labeled gene sets, we have demonstrated that ASD genes exhibit unique spatiotemporal gene expression patterns compared to non-risk genes. The gene-level constraint metric features are complementary to gene expression features. A combination of both types of features achieves the highest accuracy in predicting ASD genes. We have demonstrated the validity of our method through extensive validations using independent sets of risk genes and differential gene expression evidence. We have further shown the superior performance of our ranking system over several other state-of-the-art ranking systems.

We explored the potential role of the top ranked genes in ASD risk. The gene *HERC1*, which encodes a protein that is a probable E3 ubiquitin-protein ligase, was assigned the highest probability for conferring ASD risk. Indeed, multiple lines of evidence indicate a role for *HERC1* in ASD: 1) it was reported that *HERC1* mutations caused intellectual disability and facial dysmorphism in two Colombian siblings (37); 2) A nonsense variant in *HERC1* was associated with intellectual disability, megalencephaly, thick corpus callosum and cerebellar atrophy (38); 3) importantly, mutations in *HERC1* were reported to be associated with ASD in an exome sequencing study (39). Another notable gene in our top list was *TCF20* (ranked seventh), which encodes a transcription activator. Intriguingly, *TCF20* was one of the highest ranking candidate autism risk genes (category 2) according to the most recent version of the SFARI Gene resource. Mutations in *TCF20* were also implicated in Phelan-McDermid syndrome (40), developmental disorders (41), and schizophrenia (42). Our ranking system also successfully predicted another ASD gene *FBXO11* (ranked sixth). *FBXO11* was prioritized as a strong ASD candidate gene (43), and was recently reported to be associated with a variable neurodevelopmental disorder (44). Additionally, mutations in another two genes, *NBEA1* and *ATRX,* ranked fourth and fifth in our analysis, have also been identified with ASD according to SFARI Gene resource.

Our ranking system also highlighted some potential novel candidate genes that may deserve further investigation. Three genes, *CAND1*, *DOCK3*, and *MYCBP2,* ranked second, eighth and tenth, are all involved in neuronal development. To our knowledge, direct genetic links between these genes with ASD have not been found. *CAND1* encodes an essential regulator of Cullin-RING ubiquitin ligases that play a critical role in ubiquitination and protein degradation (45). Of note, the ubiquitination pathway has been implicated in neuronal function and brain disorders, including ASD (46). *DOCK3* encodes dedicator of cytokinesis 3, which induces axonal outgrowth in the central nervous system. *DOCK3* has been associated with Alzheimer disease (47), epilepsy (48), intellectual disability (49), and attention deficit hyperactivity disorder (50). *MYCBP2* encodes an E3 ubiquitin-protein ligase that plays a role in axon guidance and synapse formation in the developing nervous system. *MYCBP2* has been implicated in acute lymphoblastic leukemia and a rare inherited vision defect (51, 52). We have provided the whole list of ranked genes with their probability scores in Supplementary Table 2. Researchers can further explore the top-ranked genes or genes of their own interest.

Our study not only provides hundreds of new ASD candidate genes with strong evidence for involvement in ASD, but also shows that the predicted risk genes are indeed biologically meaningful and are clustered around biological processes relevant to ASD. GO enrichment analysis demonstrated that the predicted risk genes were enriched in GO terms related to neuronal signaling, neurogenesis, chromatin remodeling, and histone modification, all of which are important biological processes implicated in ASD. In addition, among our top ten ranked genes, we found that five were related to the ubiquitination pathway (*HERC1, CAND1, ZYG11B, FBXO11,* and *MYCBP2*), which is consistent with the significant enrichment of protein ubiquitination process in our GO enrichment analysis (GO_PROTEIN_UBIQUITINATION, p_corrected_ = 9 × 10^−6^), supporting the merging role of ubiquitin signaling in ASD (46, 53). Our analyses also highlighted other biological mechanisms that may underlie ASD. For example, there is evidence for roles of RNA alternative splicing (54) and RNA polyadenylation (55) in ASD, both of which were represented in our top enriched GO terms.

Our study also sheds light on when and where ASD genes may exert their effects during brain development. Of the 14 gene expression features from the top 30 important features in the random forest model, nine referred to brain regions in the early to mid-prenatal stage (≤ 16 pcw), reinforcing the important role of early prenatal development in ASD. The involved brain regions include the mediodorsal nucleus of the thalamus (MD), dorsal thalamus (DTH), ventrolateral prefrontal cortex (VFC), primary motor cortex (M1C), dorsolateral prefrontal cortex (DFC), and medial prefrontal cortex (MFC). All of these brain regions have been implicated in ASD pathology (56). Another notable gene expression feature occurred in the late prenatal stage (35 pcw) in cerebellum (CBC), supporting the emerging role of this region in ASD (57).

In summary, our study has demonstrated that human brain spatiotemporal gene expression patterns and gene-level constraint metrics predict ASD risk genes. Our gene ranking system provides a useful resource for prioritizing ASD candidate genes.

## Acknowledgments

This study was supported by National Institutes of Health grants R01 AA022994 and AA024486 (to S.H.).

---

**Supplementary Figure 1. Heatmap view of spatiotemporal gene expression in human brain.** Each cell in the heat map corresponds to the expression level of a gene (row) in a specific brain region and development stage (column). The ASD risk and non-risk genes are denoted by red and blue rows respectively. Brain regions are represented by the 31 colors in Color Key of Brain Regions. The intensity of the color in each cell represents the log2-transformed expression level.

**Supplementary Figure 2. Boxplot of gene-level constraint metrics and other gene features for true positive (TP) and true negative (TN) genes.**

**Supplementary Figure 3. Boxplot of AUCs under different feature sets for random forest model.**

**Supplementary Figure 4. Enrichment analysis of down-regulated differential expression evidence for predicted ASD risk genes.**

**Supplementary Figure 5. Hierarchical clustering of 276 significant GO terms.**

**Supplementary Figure 6. Comparison of our gene ranking system (random forest, RF) with five other gene ranking systems on classifying true positive and true negative genes.**

**Supplementary Figure 7. Comparison of our gene ranking system (random forest, RF) with five other gene ranking systems on overall rankings of 173 independent candidate genes**

**Supplementary Figure 8. Spatiotemporal gene expression patterns in human brain for the top ten ranked genes**

## Reference

1. Jeste SS, Geschwind DH. Disentangling the heterogeneity of autism spectrum disorder through genetic findings. Nature reviews Neurology. 2014;10(2):74-81.

2. Colvert E, Tick B, McEwen F, Stewart C, Curran SR, Woodhouse E, et al. Heritability of Autism Spectrum Disorder in a UK Population-Based Twin Sample. JAMA psychiatry. 2015;72(5):415-23.

3. Hallmayer J, Cleveland S, Torres A, Phillips J, Cohen B, Torigoe T, et al. Genetic heritability and shared environmental factors among twin pairs with autism. Archives of general psychiatry. 2011;68(11):1095-102.

4. Stessman HA, Xiong B, Coe BP, Wang T, Hoekzema K, Fenckova M, et al. Targeted sequencing identifies 91 neurodevelopmental-disorder risk genes with autism and developmental-disability biases. Nature genetics. 2017;49(4):515-26.

5. Wang T, Guo H, Xiong B, Stessman HA, Wu H, Coe BP, et al. De novo genic mutations among a Chinese autism spectrum disorder cohort. Nature communications. 2016;7:13316.

6. Ronemus M, Iossifov I, Levy D, Wigler M. The role of de novo mutations in the genetics of autism spectrum disorders. Nature reviews Genetics. 2014;15(2):133-41.

7. Sanders SJ, Murtha MT, Gupta AR, Murdoch JD, Raubeson MJ, Willsey AJ, et al. De novo mutations revealed by whole-exome sequencing are strongly associated with autism. Nature. 2012;485(7397):237-41.

8. Iossifov I, Ronemus M, Levy D, Wang Z, Hakker I, Rosenbaum J, et al. De novo gene disruptions in children on the autistic spectrum. Neuron. 2012;74(2):285-99.

9. Iossifov I, O’Roak BJ, Sanders SJ, Ronemus M, Krumm N, Levy D, et al. The contribution of de novo coding mutations to autism spectrum disorder. Nature. 2014;515(7526):216-21.

10. RK CY, Merico D, Bookman M, J LH, Thiruvahindrapuram B, Patel RV, et al. Whole genome sequencing resource identifies 18 new candidate genes for autism spectrum disorder. Nature neuroscience. 2017;20(4):602-11.

11. Turner TN, Coe BP, Dickel DE, Hoekzema K, Nelson BJ, Zody MC, et al. Genomic Patterns of De Novo Mutation in Simplex Autism. Cell. 2017;171(3):710-22 e12.

12. Xu J, Li Y. Discovering disease-genes by topological features in human protein-protein interaction network. Bioinformatics. 2006;22(22):2800-5.

13. Gandhi TK, Zhong J, Mathivanan S, Karthick L, Chandrika KN, Mohan SS, et al. Analysis of the human protein interactome and comparison with yeast, worm and fly interaction datasets. Nature genetics. 2006;38(3):285-93.

14. Krumm N, O’Roak BJ, Shendure J, Eichler EE. A de novo convergence of autism genetics and molecular neuroscience. Trends in neurosciences. 2014;37(2):95-105.

15. Gilman SR, Iossifov I, Levy D, Ronemus M, Wigler M, Vitkup D. Rare de novo variants associated with autism implicate a large functional network of genes involved in formation and function of synapses. Neuron. 2011;70(5):898-907.

16. Li J, Shi M, Ma Z, Zhao S, Euskirchen G, Ziskin J, et al. Integrated systems analysis reveals a molecular network underlying autism spectrum disorders. Molecular systems biology. 2014;10:774.

17. Hormozdiari F, Penn O, Borenstein E, Eichler EE. The discovery of integrated gene networks for autism and related disorders. Genome research. 2015;25(1):142-54.

18. Liu L, Lei J, Roeder K. Network Assisted Analysis to Reveal the Genetic Basis of Autism. The annals of applied statistics. 2015;9(3):1571-600.

19. Krishnan A, Zhang R, Yao V, Theesfeld CL, Wong AK, Tadych A, et al. Genome-wide prediction and functional characterization of the genetic basis of autism spectrum disorder. Nature neuroscience. 2016;19(11):1454-62.

20. Duda M, Zhang H, Li HD, Wall DP, Burmeister M, Guan Y. Brain-specific functional relationship networks inform autism spectrum disorder gene prediction. Translational psychiatry. 2018;8(1):56.

21. Willsey AJ, Sanders SJ, Li M, Dong S, Tebbenkamp AT, Muhle RA, et al. Coexpression networks implicate human midfetal deep cortical projection neurons in the pathogenesis of autism. Cell. 2013;155(5):997-1007.

22. Samocha KE, Robinson EB, Sanders SJ, Stevens C, Sabo A, McGrath LM, et al. A framework for the interpretation of de novo mutation in human disease. Nature genetics. 2014;46(9):944-50.

23. Iossifov I, Levy D, Allen J, Ye K, Ronemus M, Lee YH, et al. Low load for disruptive mutations in autism genes and their biased transmission. Proceedings of the National Academy of Sciences of the United States of America. 2015;112(41):E5600-7.

24. Petrovski S, Wang Q, Heinzen EL, Allen AS, Goldstein DB. Genic intolerance to functional variation and the interpretation of personal genomes. PLoS genetics. 2013;9(8):e1003709.

25. Lek M, Karczewski KJ, Minikel EV, Samocha KE, Banks E, Fennell T, et al. Analysis of protein-coding genetic variation in 60,706 humans. Nature. 2016;536(7616):285-91.

26. Kosmicki JA, Samocha KE, Howrigan DP, Sanders SJ, Slowikowski K, Lek M, et al. Refining the role of de novo protein-truncating variants in neurodevelopmental disorders by using population reference samples. Nature genetics. 2017;49(4):504-10.

27. Sanders SJ, He X, Willsey AJ, Ercan-Sencicek AG, Samocha KE, Cicek AE, et al. Insights into Autism Spectrum Disorder Genomic Architecture and Biology from 71 Risk Loci. Neuron. 2015;87(6):1215-33.

28. Rossin EJ, Lage K, Raychaudhuri S, Xavier RJ, Tatar D, Benita Y, et al. Proteins encoded in genomic regions associated with immune-mediated disease physically interact and suggest underlying biology. PLoS genetics. 2011;7(1):e1001273.

29. Bonacich P. Power and Centrality: A Family of Measures. American Journal of Sociology. 1987;92:1170-82.

30. Batagelj V, Zaversnik M. An O(m) Algorithm for Cores Decomposition of Networks. arXiv preprint. 2003;cs/0310049.

31. Brin S, Page L, editors. The Anatomy of a Large-Scale Hypertextual Web Search Engine. Proceedings of the 7th World-Wide Web Conference; 1998; Brisbane, Australia,.

32. Gandal MJ, Haney JR, Parikshak NN, Leppa V, Ramaswami G, Hartl C, et al. Shared molecular neuropathology across major psychiatric disorders parallels polygenic overlap. Science. 2018;359(6376):693-7.

33. Subramanian A, Tamayo P, Mootha VK, Mukherjee S, Ebert BL, Gillette MA, et al. Gene set enrichment analysis: a knowledge-based approach for interpreting genome-wide expression profiles. Proceedings of the National Academy of Sciences of the United States of America. 2005;102(43):15545-50.

34. Chen YA, Tripathi LP, Dessailly BH, Nystrom-Persson J, Ahmad S, Mizuguchi K. Integrated pathway clusters with coherent biological themes for target prioritisation. PloS one. 2014;9(6):e99030.

35. Zhang C, Shen Y. A Cell Type-Specific Expression Signature Predicts Haploinsufficient Autism-Susceptibility Genes. Human mutation. 2017;38(2):204-15.

36. Li J, Wang L, Guo H, Shi L, Zhang K, Tang M, et al. Targeted sequencing and functional analysis reveal brain-size-related genes and their networks in autism spectrum disorders. Molecular psychiatry. 2017;22(9):1282-90.

37. Ortega-Recalde O, Beltran OI, Galvez JM, Palma-Montero A, Restrepo CM, Mateus HE, et al. Biallelic HERC1 mutations in a syndromic form of overgrowth and intellectual disability. Clinical genetics. 2015;88(4):e1-3.

38. Nguyen LS, Schneider T, Rio M, Moutton S, Siquier-Pernet K, Verny F, et al. A nonsense variant in HERC1 is associated with intellectual disability, megalencephaly, thick corpus callosum and cerebellar atrophy. European journal of human genetics: EJHG. 2016;24(3):455-8.

39. Hashimoto R, Nakazawa T, Tsurusaki Y, Yasuda Y, Nagayasu K, Matsumura K, et al. Whole-exome sequencing and neurite outgrowth analysis in autism spectrum disorder. Journal of human genetics. 2016;61(3):199-206.

40. Upadia J, Gonzales PR, Atkinson TP, Schroeder HW, Robin NH, Rudy NL, et al. A previously unrecognized 22q13.2 microdeletion syndrome that encompasses TCF20 and TNFRSF13C. American journal of medical genetics Part A. 2018.

41. Deciphering Developmental Disorders S. Prevalence and architecture of de novo mutations in developmental disorders. Nature. 2017;542(7642):433-8.

42. Smeland OB, Frei O, Kauppi K, Hill WD, Li W, Wang Y, et al. Identification of Genetic Loci Jointly Influencing Schizophrenia Risk and the Cognitive Traits of Verbal-Numerical Reasoning, Reaction Time, and General Cognitive Function. JAMA psychiatry. 2017;74(10):1065-75.

43. Ji X, Kember RL, Brown CD, Bucan M. Increased burden of deleterious variants in essential genes in autism spectrum disorder. Proceedings of the National Academy of Sciences of the United States of America. 2016;113(52):15054-9.

44. Gregor A, Sadleir LG, Asadollahi R, Azzarello-Burri S, Battaglia A, Ousager LB, et al. De Novo Variants in the F-Box Protein FBXO11 in 20 Individuals with a Variable Neurodevelopmental Disorder. American journal of human genetics. 2018.

45. Zheng J, Yang X, Harrell JM, Ryzhikov S, Shim EH, Lykke-Andersen K, et al. CAND1 binds to unneddylated CUL1 and regulates the formation of SCF ubiquitin E3 ligase complex. Molecular cell. 2002;10(6):1519-26.

46. Mabb AM, Ehlers MD. Ubiquitination in postsynaptic function and plasticity. Annual review of cell and developmental biology. 2010;26:179-210.

47. Tachi N, Hashimoto Y, Matsuoka M. MOCA is an integrator of the neuronal death signals that are activated by familial Alzheimer’s disease-related mutants of amyloid beta precursor protein and presenilins. The Biochemical journal. 2012;442(2):413-22.

48. Li J, Mi X, Chen L, Jiang G, Wang N, Zhang Y, et al. Dock3 Participate in Epileptogenesis Through rac1 Pathway in Animal Models. Molecular neurobiology. 2016;53(4):2715-25.

49. Helbig KL, Mroske C, Moorthy D, Sajan SA, Velinov M. Biallelic loss-of-function variants in DOCK3 cause muscle hypotonia, ataxia, and intellectual disability. Clinical genetics. 2017;92(4):430-3.

50. de Silva MG, Elliott K, Dahl HH, Fitzpatrick E, Wilcox S, Delatycki M, et al. Disruption of a novel member of a sodium/hydrogen exchanger family and DOCK3 is associated with an attention deficit hyperactivity disorder-like phenotype. Journal of medical genetics. 2003;40(10):733-40.

51. Bredrup C, Johansson S, Bindoff LA, Sztromwasser P, Krakenes J, Mellgren AE, et al. High myopia-excavated optic disc anomaly associated with a frameshift mutation in the MYC-binding protein 2 gene (MYCBP2). American journal of ophthalmology. 2015;159(5):973-9 e2.

52. Ge Z, Guo X, Li J, Hartman M, Kawasawa YI, Dovat S, et al. Clinical significance of high c-MYC and low MYCBP2 expression and their association with Ikaros dysfunction in adult acute lymphoblastic leukemia. Oncotarget. 2015;6(39):42300-11.

53. Cheon S, Dean M, Chahrour M. The ubiquitin proteasome pathway in neuropsychiatric disorders. Neurobiology of learning and memory. 2018.

54. Parikshak NN, Swarup V, Belgard TG, Irimia M, Ramaswami G, Gandal MJ, et al. Genome-wide changes in lncRNA, splicing, and regional gene expression patterns in autism. Nature. 2016;540(7633):423-7.

55. Parras A, Anta H, Santos-Galindo M, Swarup V, Elorza A, Nieto-Gonzalez JL, et al. Autism-like phenotype and risk gene mRNA deadenylation by CPEB4 mis-splicing. Nature. 2018;560(7719):441-6.

56. Ha S, Sohn IJ, Kim N, Sim HJ, Cheon KA. Characteristics of Brains in Autism Spectrum Disorder: Structure, Function and Connectivity across the Lifespan. Experimental neurobiology. 2015;24(4):273-84.

57. Becker EB, Stoodley CJ. Autism spectrum disorder and the cerebellum. International review of neurobiology. 2013;113:1-34.

58. Potkin SG, Guffanti G, Lakatos A, Turner JA, Kruggel F, Fallon JH, et al. Hippocampal atrophy as a quantitative trait in a genome-wide association study identifying novel susceptibility genes for Alzheimer’s disease. PloS one. 2009;4(8):e6501.

59. Castermans D, Wilquet V, Parthoens E, Huysmans C, Steyaert J, Swinnen L, et al. The neurobeachin gene is disrupted by a translocation in a patient with idiopathic autism. Journal of medical genetics. 2003;40(5):352-6.

60. Castermans D, Volders K, Crepel A, Backx L, De Vos R, Freson K, et al. SCAMP5, NBEA and AMISYN: three candidate genes for autism involved in secretion of large dense-core vesicles. Human molecular genetics. 2010;19(7):1368-78.

61. Gong X, Bacchelli E, Blasi F, Toma C, Betancur C, Chaste P, et al. Analysis of X chromosome inactivation in autism spectrum disorders. American journal of medical genetics Part B, Neuropsychiatric genetics: the official publication of the International Society of Psychiatric Genetics. 2008;147B(6):830-5.

62. Watson LA, Goldberg H, Berube NG. Emerging roles of ATRX in cancer. Epigenomics. 2015;7(8):1365-78.

63. Redin C, Gerard B, Lauer J, Herenger Y, Muller J, Quartier A, et al. Efficient strategy for the molecular diagnosis of intellectual disability using targeted high-throughput sequencing. Journal of medical genetics. 2014;51(11):724-36.

64. Hardisty-Hughes RE, Tateossian H, Morse SA, Romero MR, Middleton A, Tymowska-Lalanne Z, et al. A mutation in the F-box gene, Fbxo11, causes otitis media in the Jeff mouse. Human molecular genetics. 2006;15(22):3273-9.

65. Fritzen D, Kuechler A, Grimmel M, Becker J, Peters S, Sturm M, et al. De novo FBXO11 mutations are associated with intellectual disability and behavioural anomalies. Human genetics. 2018;137(5):401-11.

66. Gregor A, Sadleir LG, Asadollahi R, Azzarello-Burri S, Battaglia A, Ousager LB, et al. De Novo Variants in the F-Box Protein FBXO11 in 20 Individuals with a Variable Neurodevelopmental Disorder. American journal of human genetics. 2018;103(2):305-16.

67. Babbs C, Lloyd D, Pagnamenta AT, Twigg SR, Green J, McGowan SJ, et al. De novo and rare inherited mutations implicate the transcriptional coregulator TCF20/SPBP in autism spectrum disorder. Journal of medical genetics. 2014;51(11):737-47.

68. Lelieveld SH, Reijnders MR, Pfundt R, Yntema HG, Kamsteeg EJ, de Vries P, et al. Meta-analysis of 2,104 trios provides support for 10 new genes for intellectual disability. Nature neuroscience. 2016;19(9):1194-6.

69. Iwata-Otsubo A, Ritter AL, Weckselbatt B, Ryan NR, Burgess D, Conlin LK, et al. DOCK3-related neurodevelopmental syndrome: Biallelic intragenic deletion of DOCK3 in a boy with developmental delay and hypotonia. American journal of medical genetics Part A. 2018;176(1):241-5.

70. Erdem-Eraslan L, Heijsman D, de Wit M, Kremer A, Sacchetti A, van der Spek PJ, et al. Tumor-specific mutations in low-frequency genes affect their functional properties. Journal of neuro-oncology. 2015;122(3):461-70.

